# Reproducible-by-design: Romics Processor, a FAIR ecosystem for multi-omics and spatial-omics analysis

**DOI:** 10.64898/2026.07.09.737600

**Authors:** Brittney L. Gorman, Harsh Bhotika, Matthew Jehrio, Jeffrey M. Purkerson, Frederic Carlin, Ernesto Nakayasu, Andrew M. Dylag, Ravi S. Misra, Joshua Adkins, Christopher R. Anderton, Gloria Pryhuber, Geremy Clair

## Abstract

Multi-omics and spatial-omics technologies are exploding in use, producing increasingly complex datasets. Existing bioinformatics tools are developing rapidly but fail to fully enforce the FAIR principles, leaving the field vulnerable to escalating issues in computational reproducibility. Here, we introduce a reproducible-by-design paradigm represented in an omics data processing package, RomicsProcessor. At its core, the “Romics_object”, which is a self-contained digital artifact that encapsulates the full history of the data from the original data to the fully processed state, capturing the details of the transformative steps and the required dependencies. This architecture ensures that computational workflows are fully portable and reproducible. In this manuscript, we demonstrate RomicProcessor’s computational capabilities and scalability on diverse datasets, including bulk proteomics, large-scale multiplexed immunofluorescence, and multi-batch mass spectrometry imaging. Providing a robust framework for truly FAIR Data Principles-based analysis, RomicsProcessor is a blueprint for the next generation of reproducible bioinformatics tools that can dramatically accelerate discovery in multi-omics biology in the era of artificial intelligence.

## Introduction

The widespread adoption of high-throughput omics technologies has created a data analysis reproducibility crisis^1–3^. While sharing raw data and data analysis scripts is now a common practice, these efforts are fundamentally insufficient to guarantee reproducibility. As widely recognized by both funding agencies and researchers, the script-and-data model of sharing is inherently fragile, often collapsing due to various factors^2,4–8^. This problem is further complicated as the omics fields expand towards complex spatial methods such as spatial transcriptomics, highly multiplexed immunofluorescence, and mass spectrometry imaging^5,9–11^. To overcome these challenges, a set of guiding principles was established in 2016: the FAIR principles (Findability, Accessibility, Interoperability, and Reusability)^12^, yet their full implementation within software design remains limited.

In practice, this fragility manifests in several ways. Despite widespread adoption of raw data sharing frameworks across omics communities, approaches for transparently sharing data processing steps remain insufficient^13,14^. While some researchers now share their analysis scripts, such as R markdowns and Python Jupyter notebooks^15,16^, these efforts often fail to guarantee end-to-end reproducibility. Key processing steps are sometimes omitted from published reports, or proprietary processing tools are used ‘in the loop’, hindering reproducible data analysis and data reuse. These limitations arise in part from unrecorded dependencies on specific software packages and versions, as well as omission of essential contextual metadata and finely tuned parameters from final reports.

To overcome these fundamental limitations, we propose a shift in computational analysis: moving from separate, disconnected components to a single, self-contained analytical object. Our approach to create this “reproducibility-by-design” object is to ensure that such object contains the data’s entire lifecycle: from its original state through every transformation to the published results, along with the complete metadata and details of the computational environment.

RomicsProcessor is designed to address reproducibility challenges by offering a robust, transparent framework for analyzing bulk, single-cell, and spatial omics datasets. Supporting various data types, our framework provides a comprehensive set of functionalities for end-to-end analysis including data import, transformation, normalization, subsampling, dimensionality reduction, clustering, statistical analysis, metadata operations and visualization tools for quality control and data exploration. At the core of our framework, the “Romics_object” acts as a fully traceable, self-contained digital artifact of computational history (Fig. 1A). RomicsProcessor automatically logs all processing steps (including user-defined parameters) and dependencies (including package versions) alongside the original_data, original_metadata, and their processed versions. By doing so, it guarantees that an analysis can be identically replicated from the details contained in an object alone, obviating the fragility of external scripts and manual record-keeping. This approach provides reproducibility at the analytical logic layer, complementing system-level containerization tools such as Docker that ensure reproducibility at the operating system level^17,18^. Importantly, RomicsProcessor allows the creation of full processing pipelines with a single line of code. Such pipelines can be applied for reproducibility verification or to other objects, facilitating streamlined and reproducible data analysis.

**Figure 1:**
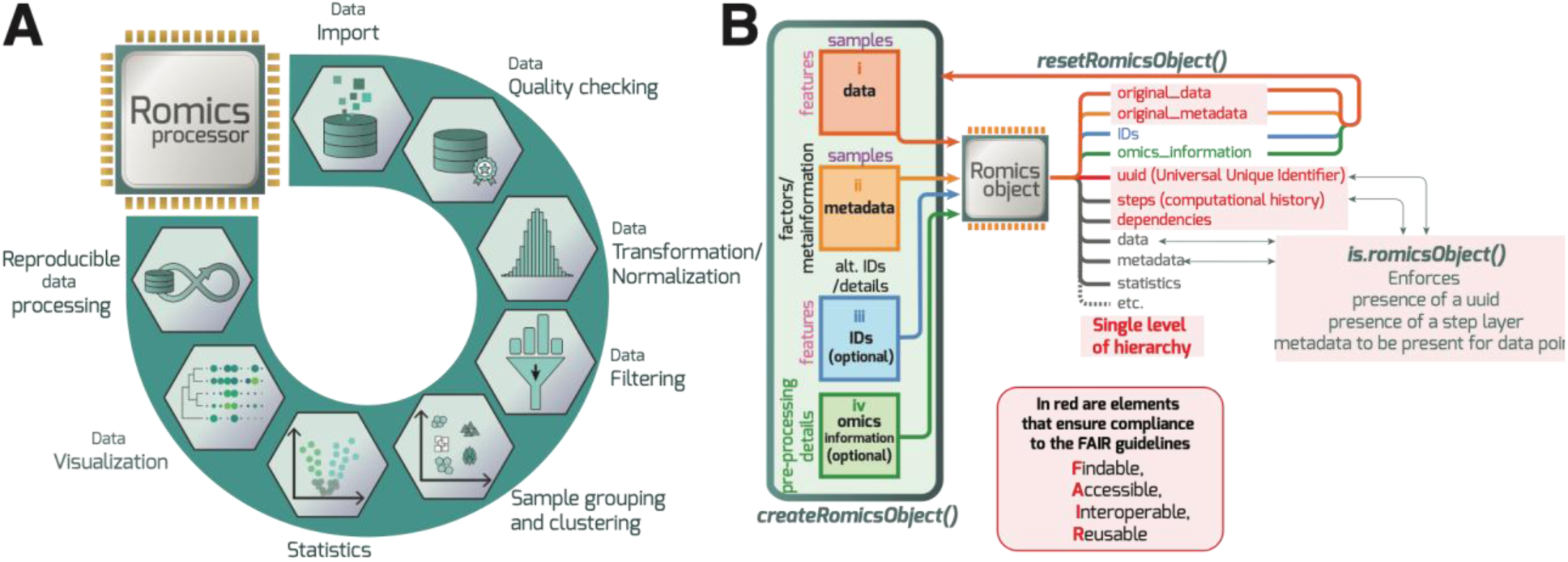
RomicsProcessor an example of FAIR-compatible package implementation. **A.** RomicsProcessor is an all-in-one solution for omics data processing including modules to perform various steps of data processing, statistics, and visualization. **B.** The Romics_object structure is designed to store a universal unique identifier, original data, metadata, the transformation steps taken with the data, and package dependency versions. The *is.romicsObject()* function checks whether the object remains compatible with FAIR practices.

In this manuscript, we present RomicsProcessor not only as an omics analysis ecosystem, but as a generalizable blueprint for developing FAIR-native computational tools. We illustrate through three use cases how RomicsProcessor can be utilized to accelerate discoveries from omics and spatial omics datasets. Finally, we provide a standard for implementing an end-to-end object-centric architecture, paving the way for a new generation of sustainable, replicable analysis solutions across programing languages and scientific domains.

## Results

### Playbook for a FAIR-by-design R package: RomicsProcessor

The core of RomicsProcessor’s FAIR compliance is its reproducible-by-design object architecture. We designed the “Romics_object” (Fig. 1B) to programmatically enforce FAIR principles^12^ through its structure, integrity checks and a suite of dedicated functions. This section describes the playbook for these design choices.

The input required for Romics_objects are standard tables familiar to users of other processing packages^19,20^: (i) a “data” table, where the columns represent the samples (or spatial pixels), and the rows the analytical “features” (e.g., transcripts, proteins, lipids, metabolites) and (ii) a “metadata” table, with the columns representing samples (or spatial pixels) and the rows representing the metadata “factors” (or metainformation). A key control step during the object creation is that the data and metadata must have the same number of columns, named identically. For spatial datasets, the x, y, and z coordinates of the samples can be stored directly in the metadata. Two optional files can also be imported: (iii) an “IDs” table containing alternative IDs or feature details, and (iv) an “omics_information” table. We recommend using this layer to record pre-processing details that occur prior to import, such as parameters from preprocessing mass spectrometry or imaging analysis software programs.

<u>F</u>indability: To ensure every dataset is uniquely findable, a Universally Unique Identifier (UUID) is automatically generated and embedded upon creation in the Romics_object. This UUID can be used to track objects that are derivative or branches from the original object, guaranteeing analytical traceability. Transformative functions return a new Romics_object derived from the previous one using the syntax “object_B <-*transformativeFunction*(object_A)”. The function *checkRelationRomicsObjects()* determines if two objects are derived from one another and indicates the branching point and differences between them. In addition, the objects store the “original_data” and “original_metadata” tables allowing any analysis to be traced back to its starting point. To preserve this original state, the package intentionally provides no function that can modify these layers. The function *resetRomicsObject()* restores the objects to their initial form, allowing for reproducibility testing.

<u>A</u>ccessibility: The Romics_object’s design as a single-level list, with components like “data” and “metadata” directly accessible by name, ensures straightforward data accessibility without navigating complex hierarchies. Dedicated functions enable the retrieval of the samples, features, and factors stored in the object (e.g., *romicsSampleNames()*, *romicsFeatureNames(), romicsFactorNames()*). For complex metadata schemes, a new factor can be derived from combinations of multiple factors keeping the single level of hierarchy intact.

Interoperability: We designed the object as a single-level R list. This flat hierarchy, with explicitly named data-frames and vectors (e.g., data, original_data, steps) making the object both human- and machine-readable. This simple structure is so effective that coding assistants can readily interpret it to generate new, FAIR compliant functions when provided with RomicsProcessor source code. Interoperability is also ensured through the incorporation of helper functions to extract data directly from pre-processing tools (e.g., Fragpipe^21^, MaxQuant^22^, SCiLs^23^) and conversion functions that allow seamless transformation to and from other popular objects such as Seurat^19^ and Bioconductor SpatialExperiment^20^ objects.

<u>R</u>eusability: True reusability is a cornerstone of our design with the Romics_object capturing its own computational history. Romics_objects, contain two critical layers for this: a “dependencies” layer that records all packages names and versions used in the session, and a “steps” layer that logs every transformative function applied, including user-defined parameters and timestamps. This is implemented by a lightweight two-line code needed during the function creation. An example of RomicsProcessor function implementation is provided as in the Supplementary material. The structural integrity of the object and its FAIR compliance can be enforced by an internal *is.romicsObject()* function that systematically validates the presence and integrity of these crucial layers along with the “original_data” and “original_metadata” object.

The culmination of the framework is the *createRomicsPipeline()* function which converts the object’s stored history back to executable R code that can be applied to other Romics_objects in a single command. Thus, it can be applied to the original object for a one command reproducibility verification or to new datasets for consistent, high-throughput processing.

### RomicsProcessor: a versatile multi-omics processor tool

Though not previously published as a method, the versatility and robustness of RomicsProcessor are demonstrated by its use in peer reviewed research. It has been instrumental in a variety of omics studies using diverse technologies including bulk proteomics^24–26^, lipidomics^27^, spatially resolved proteomics^28^ and transcriptomics^29^, as well as single-cell proteomics^30,31^. This proven track record is built upon a comprehensive suite of features covering the entire analytical pipeline, from data import and pre-processing (e.g., missing values handling, transformation, normalization) to core analysis, statistical testing, and the generation of publication-ready visualizations (e.g., UMAP, volcano plots, violin plots, heatmaps, spatial maps, see Fig. 1A). In the following use cases, we further demonstrate RomicsProcessor scalability to large and complex datasets from immunofluorescence and mass spectrometry imaging.

Like many other groups^8,12,32,33^, we experience firsthand how cumbersome it is to ensure end-to-end reproducibility. Adhering to the FAIR principles often requires sharing raw data, metadata, code, and environment details separately, creating a fragile system where everything must be properly linked and cross-referenced. This challenge motivated us to progressively evolve RomicsProcessor, designing the Romics_object as a self-contained digital artifact that guarantees analytical reproducibility. Notably, the self-contained nature of these objects also simplifies the deployment of interactive Shiny web interfaces, enabling data exploration by collaborators without coding expertise. The following three use cases demonstrate these capabilities in action.

### Case study 1: Robust pipeline generation and reproducible omics analysis using RomicsProcessor

Our first case study demonstrates RomicsProcessor’s end-to-end workflow, for reproducible analysis and biological discovery (Fig. 2A). We analyzed the proteome of the spore-forming foodborne pathogen *Bacillus cereus* grown in different media (soil extract, zucchini puree, Luria-Bertani, and a limited media AOAC, aimed to mimic the gut-environment). Detailed methods are provided in the “Supplementary methods” section. After initial pre-processing of the mass spectrometry files with FragPipe^21^, the data was imported into a Romics_object using dedicated helper functions. The log of the preprocessing setting in FragPipe (e.g., fragpipe.workflow file) was imported into the “omics_information” layer to log the upstream steps. The raw datasets were deposited in the public repository Massive under the identifier MSV000085696 (FTP at the following address: ftp://massive.ucsd.edu/MSV000085696/) ensuring end-to-end reproducibility.

**Figure 2.**
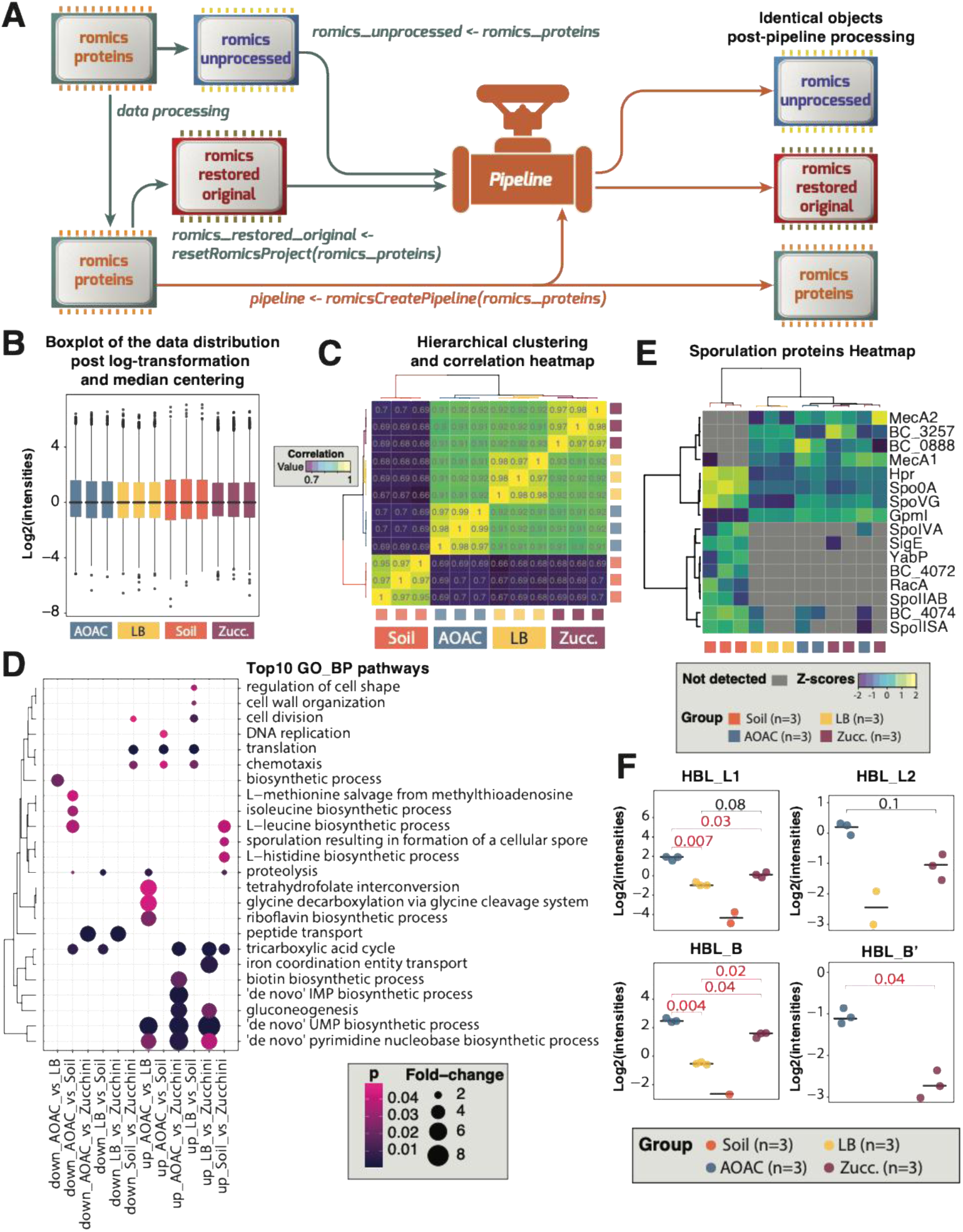
RomicsProcessor can be used to generate reproductible complex analytical pipelines. **A.** A fully processed Romics_object was used to generate a pipeline that was applied to the unprocessed and the reinitialized object resulting in identical final objects. **B.** Boxplot depicting the proteomics profile of *B. cereus* samples grown in different culture media after log2-transformation and median centering. **C.** Correlation heatmap showing that the samples grown in the same media are highly correlated (Pearson correlation > 0.95). **D.** top 10 GO terms enriched in the list of proteins higher or lower in abundance or presence in the different media comparisons (Student’s T-test or GLM binomial test adjusted-*P<0.05*, respectively). **E.** Heatmap depicting the sporulation-related proteins in the different growth media groups. F. plot showing the change of abundance of the Hbl toxin in the different growth media groups.

To test the RomicsProcessor’s reproducibility, the initial “romics_proteins” object was immediately duplicated at creation into a “romics_unprocessed” object for a downstream reproducibility challenge. Meanwhile, the “romics_proteins” object underwent a proteomics workflow including quality filtering, log-transformation, and median centering (Fig. 2B). All steps, parameters, and timestamps were automatically captured in the object’s “steps” layer. A user may choose a different workflow from the same object, yet, as long as RomicsProcessor functions are used all the steps will be recorded in the resulting object.

Quality control confirmed high within-group correlation (Pearson correlation >0.95) and clear separation between the groups by hierarchical clustering (Fig. 2C). Statistical analysis, stored in the romics_object “statistics” layer, revealed clear shifts in proteomes based on the nutrient environment. The soil extract media increased the abundance of proteins related to “sporulation” (GO 0030435, Fig. 2D-E), while the gut-mimicking media (e.g., AOAC) promoted the production of key toxins, including the components of the Hemolysin BL and the enterotoxin Nhe (Fig. 2F). This demonstrates that RomicsProcessor can be used to derive meaningful insights into bacterial pathogenesis, as both the soil-induced sporulation and specific nutrient pressure inducing enterotoxin production have previously been documented^34–36^.

Having completed the analysis, we validated the platform’s reproducibility features. Using the final processed “romics_proteins” object, a single command, *createRomicsPipeline(romics_proteins)*, generated a pipeline that could be applied to both the “romics_unprocessed” object and to a “romics_restored_original” object (created via the *resetRomicsObject()* function). In both cases, applying the pipeline resulted in identical transformed data and statistical tables, computational demonstration of RomicsProcessor’s end-to-end reproducibility (Fig. 2A). A Shiny web interface dedicated to the exploration of bulk omics dataset, RomicsExplorer, was created and is available for non-coders to visualize results contained in Romics_objects along with pre-computed statistical analyses.

### Case Study 2: Pseudo-bulk analysis of histology-guided MALDI-MSI

Our second case study demonstrates RomicsProcessor’s utility for analyzing matrix-assisted laser desorption/ionization mass spectrometry imaging (MALDI-MSI) data. We analyzed a histology-guided MALDI-MSI dataset of kidney biopsies from quality control (QC) and acute kidney injury (AKI) tissues. The data underwent histology-based segmentation and peak annotation in SCiLS, a Bruker software enabling to annotate spatial mass spectrometry imaging datasets. The data was then imported to R using a dedicated RomicsProcessor function: *extractScils()*. This function imports selected peak lists, peak alternative IDs, and region-based annotations.

After pre-processing to remove off-tissue regions, log-transformation, and median center were applied. Because the dataset spanned multiple MALDI-MSI runs, batch correction was essential to minimize batch-related signal variations. Using the QC tissue sections as a reference, we applied a batch correction, which effectively harmonized the data and removed batch effects as confirmed by a UMAP (Fig. 3A). Unsupervised Leiden clustering on the corrected data revealed 10 distinct chemical-composition-driven regions, providing a more granular map than the original histology-based segmentation of glomeruli and tubular interstitium (Fig. 3B).

**Figure 3.**
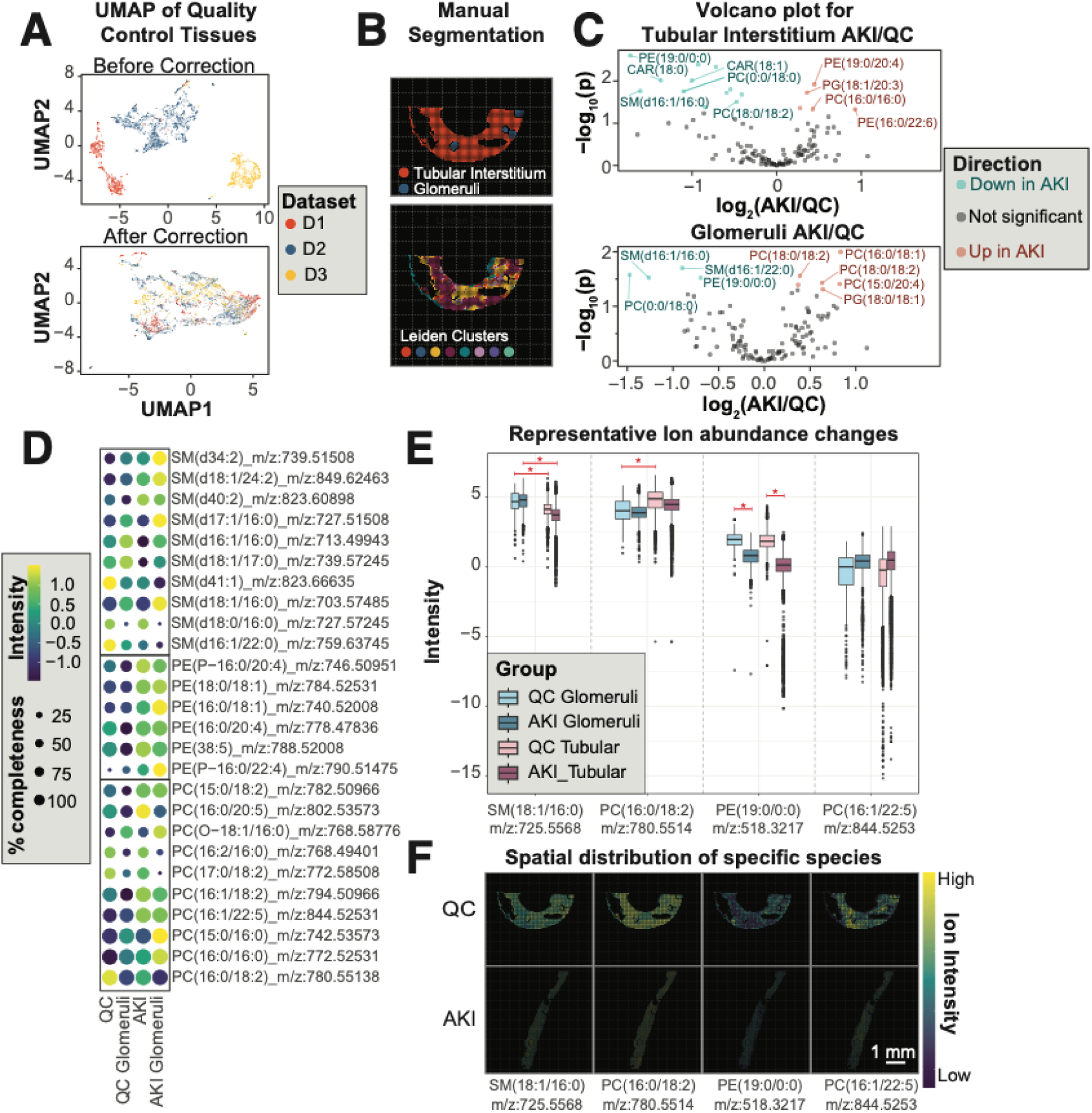
RomicsProcessor analysis of spatial-omics datasets. **A.** Perform batch correction using a quality control tissue, placed on each slide. **B.** Manual histological segmentation compared with chemical segmentation-based Leiden clustering of the lipid distributions. **C.** Volcano plots showing the pseudo-bulk Student’s T-test results (p ≤ 0.05). **D.** Bubble plot showing the most significantly changing lipids between the QC and AKI tissues (Fold-change ≥ 1). **E.** Representative boxplots of abundant ions within the glomeruli, tubules, and QC tissues (\**P* ≤ 0.05). **F.** Representative ion images of each ion shown in part (E).

To enable robust statistical comparisons, RomicsProcessor contains a suite of statistical tests (e.g., linear mixed-effect model (LMM), Pseudo-bulk transformation, ANOVA, T-tests, Generalized Linear Model (GLM) binomial tests). Here, we opted for a pseudo-bulk analysis. RomicsProcessor was used to aggregate the pixel-level data into average abundances for each histological region per sample. This approach avoids statistical inflation due to repeated pixel-level comparisons and allows straightforward application of Student’s T-tests. Our analysis recapitulated known biology, identifying established glomerular and tubular lipid markers (Fig. 3C). For instance, the glomeruli of both the AKI and QC contain elevated abundances of SM(18:1/16:0), which is a known glomerular lipid biomarker^37^, whereas the tubular interstitium contained elevated PC(16:0/18:2), a previously reported proximal tubule marker^38^. The analysis also revealed significant lipid alteration in the AKI samples (Fig. 3E), for example, a decrease in PE(19:0/0:0) in the glomeruli specifically. Importantly, RomicsProcessor’s object structure and functionality support storing pixel locations in the metadata layer alongside their intensities in the data layer, enabling direct visualization of ion images in R (Fig. 3F).

This case study demonstrates RomicsProcessor’s capacity to manage complex multi-run spatial metabolomics projects with the integration of histology-guided segmentation, handling batch corrections, and pseudo-bulk statistical workflows.

### Case study 3: Cell type identification and statistical analysis in large-scale spatial proteomics

Our last case study showcases RomicsProcessor’s versatility as it empowers the analysis of large-scale spatial proteomics data. We analyzed a 15-core human lung tissue microarray (TMA) from 6 donors across three age groups (e.g., <30, 30-40, and >50, TMA design shown in Sup. Fig. 1A) profiled with a 50-probe multiplexed immunofluorescence (MxIF) panel. This panel comprised cell boundary markers (e.g., ATP1A1, CDH1), extracellular matrix proteins (e.g., COL1A1, COL4A1), phenotypic markers (e.g., Ki67), and pulmonary cell type markers (e.g., AGER, HPGD). After segmentation with DeepCell^39^, the data and metadata were imported into a Romics_object containing over 317,000 cells.

Following import and preprocessing in RomicsProcessor (Fig. 4A-B), including log-transformation, normalization, signal thresholding to remove analytical noise, and fluorescent channel deletion (e.g., for channels that could not be denoised), a dimensionality reduction (UMAP) and Leiden clustering were performed, resolving 29 clusters (as shown on Sup. Fig1.B). The initial clustering was refined by combining three indistinct groups (Clusters 17, 18, and 28) and by sub-clustering one heterogeneous cluster (Cluster 09) that seemed to represent multiple cell populations (Sup. Fig. 1B-F). The final 30 clusters were named based on marker expression and spatial location (Fig. 4C-E). After excluding one core with emphysema pathology and the presence of patches of immune cells and erythrocytes (Sup. Fig. 1G-H), we proceeded with statistical analysis.

**Figure 4.**
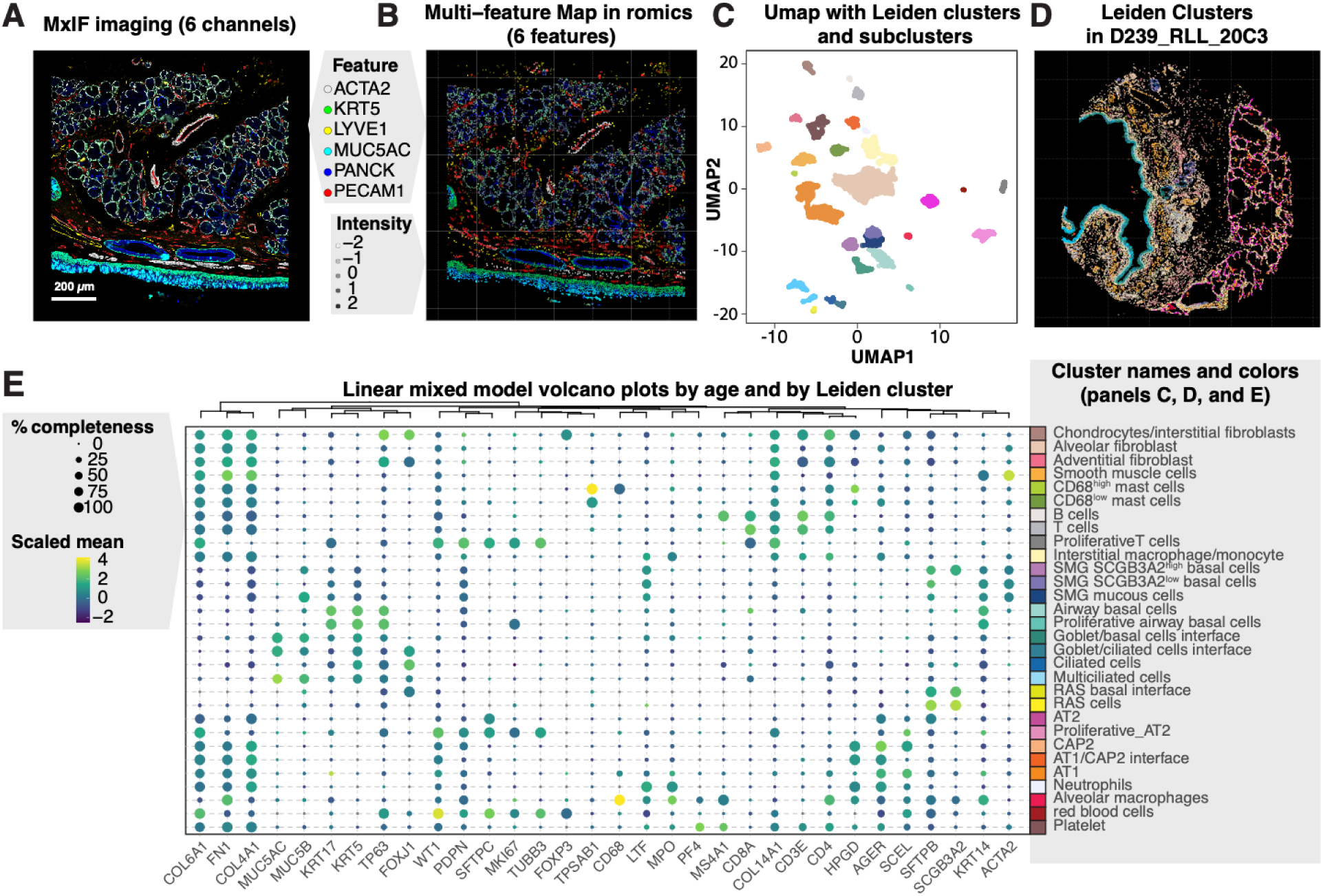
RomicsProcessor was used to process multiplexed immunofluorescence datasets. **A.** Immunofluorescence image generated using a panel of 50 antibody probes, 6 are shown here. **B.** After segmentation in DeepCell the dataset was imported in RomicsProcessor, the signal was log-transformed, normalized, and thresholded to remove analytical noise. **C.** UMAP generated from all the cells in the Romics_object colored based on the 30 Leiden clusters and subclusters. **D.** One of the biopsy cores from the TMA analyzed where each cell was colored based on cluster membership. **E.** Bubble plot showing protein abundance and percentage of detection for each of the 30 Clusters and subclusters.

As RomicsProcessor includes multiple options for computing statistics, here we opted for a different strategy than the one used in our previous case study with the use of a linear mixed-effects (LMM) test. This test is useful for capturing subtle biological signals in complex datasets while removing random effects^40^. To identify age-related protein changes while accounting for donor-to-donor variability, we leveraged the *romicsLMM()* function using the factors contained in the Romics_object as fixed and random effect. By fitting the model with the “age” as a fixed effect and the “donor” as a random effect, we discovered a significant age-associated increase of extracellular matrix proteins (e.g., COL4A1, FN1, COL14A1, EMILIN1) around the cells involved in the gas-exchange interface, namely the alveolar type 1 pneumocytes (AT1) and type 2 capillary cells (CAP2) (Fig. 5 A). This finding suggests a potential change in the composition of the fine basement layer that could reduce gas-exchange efficiency with age and highlights RomicsProcessor ability to extract subtle biological signals from complex data.

**Figure 5.**
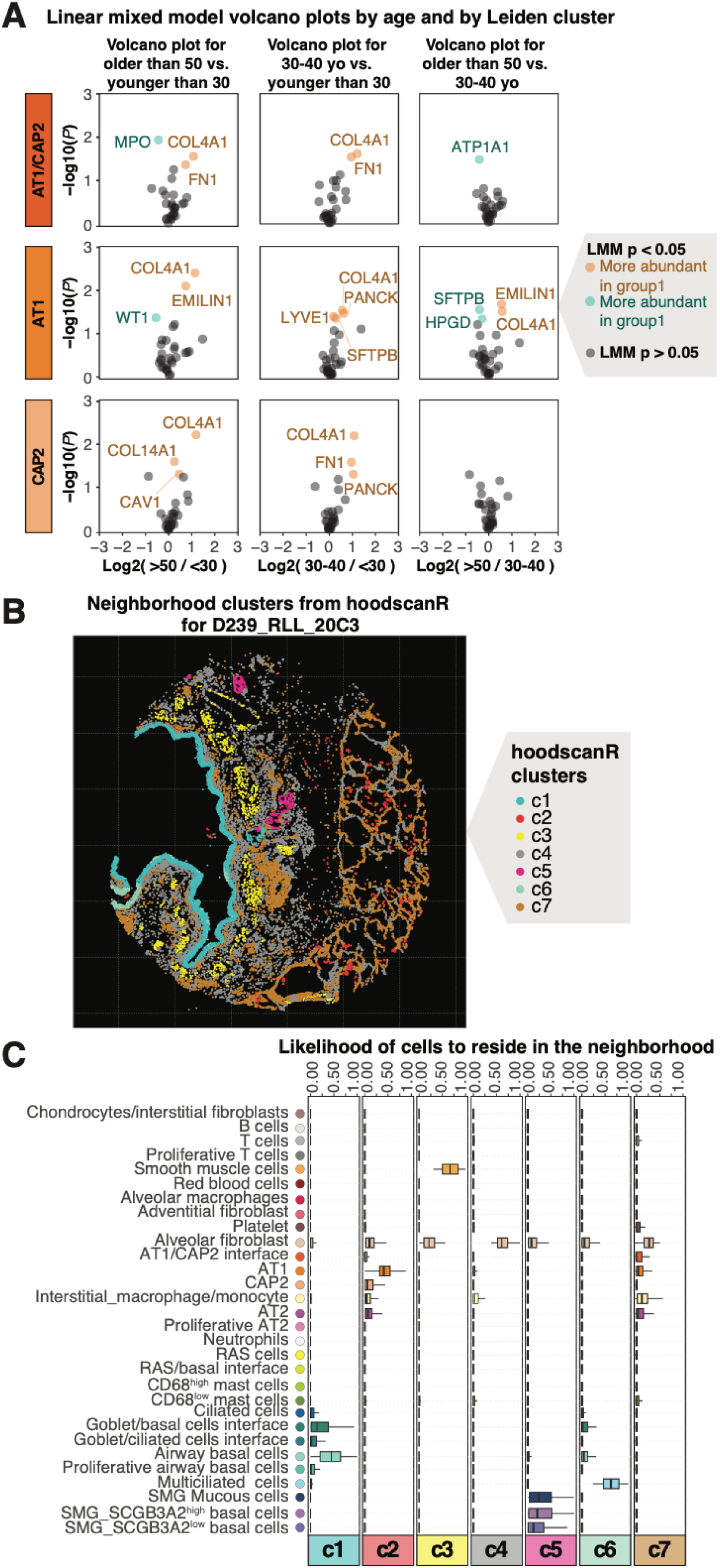
Statistical modeling and interfacing with SpatialExperiment tools. **A,** Linear mixed model Volcano plots showing ECM protein increase in the vicinity of AT1 and CAP2. **B,** hoodscanR neigborhood map and **C,** cell likelihood to reside in specific neghborhoods.

**Supplementary Figure 1.**
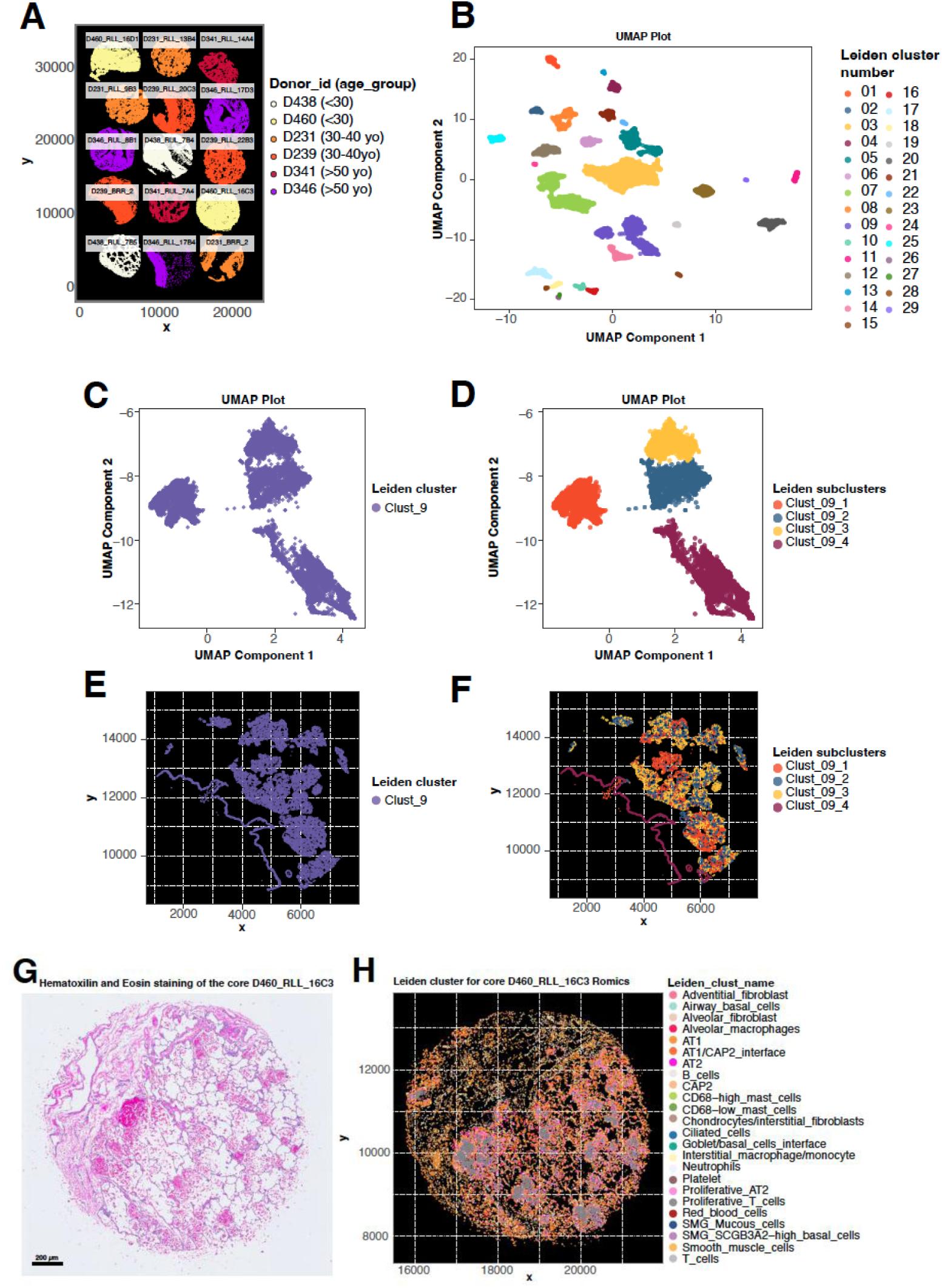
TMA design, reclustering of Leiden_cluster_OG, and tissue core removed. **A.** TMA design with donor IDs, and age group. **B.** UMAP showing the original 29 clusters. **C.** Leiden_cluster_09 in the UMAP space before re-clustering, **D.** Leiden_cluster_09 in the UMAP space after re-clustering, **E.** Leiden_cluster_09 cells before re-clustering shown on one of the TMA core coordinates. **F.** Leiden_cluster_09 cells after re-clustering, shown on one of the TMA core coordinates. **G.** and **H.** Eosin and Hematoxylin staining and Cluster position in the core D460_RLL_16C3 showing signs of emphysema pathology and immune infiltration.

Finally, to demonstrate the platform’s interoperability with the broader Bioconductor ecosystem, we converted the Romics_object to a spatialExperiment with the built-in *romicsToSpatialExperiment()* function. This enabled downstream neighborhood analysis with the external hoodscanR (v1.8.0) package, which successfully identified seven distinct tissue microenvironments (Fig. 5 B-C). Most of these corresponded to well described tissues units such as the alveolar parenchyma (neighborhood c2 and c7), submucosal glands (c5), the stratified large airway epithelium (c1 and c6), the airway interstitium (c4), or the airway smooth muscle bundles (c3). This case study establishes RomicsProcessor as a robust platform for spatial omics, validating its ability to scale to hundreds of thousands of cells, perform sophisticated statistical modeling, and seamlessly interface with other community-standard tools.

## Discussion

The implementation of RomicsProcessor fundamentally addresses the reproducibility challenges inherent to multi-omics and spatial-omics research. By moving beyond the simple sharing of scripts and data, we have introduced a new paradigm where the entire data “biography” is captured within a single, self-contained object. The elegance of this approach lies in the encapsulation of not only the processed data, but also the pristine original_data, original_metadata, and the machine-readable log of all transformative steps with the parameters and a precise record of packages dependencies. This comprehensive data structure ensures that the analysis is not only merely described in a manuscript or digital notebook but is fully and computationally reproducible, directly tackling a core bottleneck of modern data intensive biology.

A key strength of our package is its pragmatic design, promoting FAIR principle without isolating the user from the broader informatic ecosystems. It offers a comprehensive suite of internal functions but does not lock the users into a closed system. RomicsProcessor also includes converters from and to widely adopted packages such as Seurat^19^ and SpatialExperiment^20^ objects as a deliberate choice to foster interoperability. This allows researchers to leverage the unique FAIR-compliant data-handling and pipeline-generation capabilities of RomicsProcessor while still using specialized downstream tools for specific applications, as multiple functionalities remain to be implemented in the Romics ecosystem.

While we have demonstrated the package’s ability to handle large datasets, such as over 300,000 cells in the multiplex immunofluorescence (case study 3), we acknowledge that the scale of single-cell and spatial omics datasets is continually increasing, with many datasets comprising several million data points. Further testing will be required to optimize performance and memory management for these extreme scales, especially in the R language, which was not designed to be highly scalable.

However, this manuscript encourages other package developers to adopt the architectural philosophy of RomicsProcessor, as it is language-agnostic. We envision this as a “playbook” for FAIR object design and could be adopted by developers of Python-based tools such as Scanpy^41^ and Squidpy^42^. The creation of a similar PyOmicsProcessor built around the AnnData structure^43^, for example, could bring the same level of enforced reproducibility to the Python-omics community.

We recognize that Romics_objects currently only captures the analytical history after its creation. While the omics_information layer was designed to place a manual record of the crucial pre-processing details, its effectiveness relies on the diligence and intention of the data generators. To achieve true end-to-end FAIR compliance, a more enforced solution is needed. The development of universal, machine-readable “steps layer” format could ensure the seamless and verifiable transfer of information across every stage of the data life cycle.

Ultimately, the adoption of FAIR practices in data analysis^12^ could revolutionize how scientific findings are validated and used. Our implementation of RomicsProcessor enables researchers to download data objects, install the package, and execute the entire analysis in a single command. Furthermore, as AI models become more efficient at processing structured data, self-contained objects are ideal substrates for AI-driven discoveries ensuring traceability of the steps taken. We envision a future where researchers can use large language models to query these objects, re-analyzing complex datasets and testing new hypotheses in ways not originally envisioned by the data creators, truly unlocking the potential for unanticipated discoveries.

## Acknowledgements

This work was supported by the National Heart, Lung, and Blood Institute (NHLBI) Molecular Atlas of Lung Development Program Human Tissue Core (LungMAP HTC) and LungMAP BioRepository for INvestigation of Diseases of the Lung (BRINDL) through grants U01HL122700 and U01HL148861 (to GSP); U01HL122703, U01HL148860, U01HL175451 (to GC) and by the NIH Common Fund grant U54HL165443 (to GSP, with Co-Investigators GC, and CA). The work was also supported by a National Institute of Diabetes and Digestive and Kidney Diseases (NIDDK) Kidney Precision Medicine Project (KPMP) grant U01DK114920 (to CA). Part of this work was performed in the Environmental Molecular Science Laboratory, a U.S. Department of Energy (DOE) national scientific user facility at Pacific Northwest National Laboratory (PNNL). Battelle operates PNNL for the DOE under contract DE-AC05– 76RLO01830. The opinions expressed in this article are the authors’ own and do not reflect the view of the NIH, the Department of Health and Human Services, or the U.S. government. We thank Benedicte Doublet, Claire Dargaignaratz for preparing the *B. cereus* samples and Samuel Payne for sharing the raw files of the analyzed *B. cereus* samples. We are very grateful for the generosity of the donor families and honor their loss.

## Code Availability

https://github.com/PNNL-Comp-Mass-Spec/RomicsProcessor

