## Supplemental Material for "Reproducible-by-design: Romics Processor, a FAIR ecosystem for multi-omics and spatial-omics analysis"

### Supplementary methods Discussion

#### Example of RomicsProcessor straightforward function implementation.

Below is an example of implementation for a RomicsProcessor function. This simple function *medianCenterSample()* was designed to center the intensities of each sample based on their median. Lines starting by "#" are comments lines. The two lines of code in red suffice to record how the romics_object was transformed, including the user specified parameters, and must be included in any function transforming a Romics_object.

```
#The function is declared

medianCenterSample<-function(romics_object){

#This line records the user-imputed parameters

arguments<-as.list(match.call())

#This verifies that the object is a romics_object and enforces the FAIR
practices layers.

if(!is.romicsObject(romics_object) | missing(romics_object))
{stop("romics_object is missing or is not in the appropriate format")}

#This line identifies the median for each sample

col_median<- apply(romics_object$data,2,function(x) 43)
```

```
#The internal function verifies if the object was log transformed from the
step layer applies the median normalization appropriately based on this
information.

f<-function(row){

    if(romicsLogCheck(romics_object)==TRUE){

      row<-row-col_median}else{

        row<-row/col_median}

    return(row)}

#The internal function f is applied

  romics_object$data<- as.data.frame(t(apply(romics_object$data,1,f)))

#The steps are updated using the recorded user arguments

  romics_object<-romicsUpdateSteps(romics_object,arguments)

#And finally, the object is returned

  return(romics_object)

}
```

### Supplementary methods

#### *Bacillus cereus* growth and harvesting.

For the first use case, we use *Bacillus cereus* strain ATCC14579, from the INRAE SQPOV strain collection stored at -80 °C.

LB made of tryptone USP (Biokar, Beauvais, France) 10 g/L, yeast extract (Biokar) 5 g/L and NaCl (Sigma Aldrich, Saint-Louis, MO) 10 g/l. The modified AOAC (mAOAC) broth was prepared using a base of chemically defined AOAC medium M334 "Wright and Mundy (HiMedia, Mumbai, India) M334 supplemented post autoclaving with filter sterilized solutions to final concentration of 5 mM $KH_2PO_4$, 0.125 mM $MgSO_4$-$7H_2O$, 0.003 mM $MnSO_4$, 0.013 mM $ZnSO_4$-$7H_2O$, 0.02 mM iron ammonium citrate, 0.25 mM $CaCl_2$, and 6.9 mM glucose. The Zucchini Broth was prepared by steam-cooking 1 kg of fresh, washed and cut Zucchini for 10 min in a pressure cooker, after which the cooked zucchini was blended in a Stomacher at medium speed for 1 min in a Stomacher bag with lateral filter. The filtrate was centrifuged at 7,000 x g for 10 minutes and the supernatant was tyndallized by heating at 80 °C for one hour, on three consecutive days, followed by a final centrifugation step before being aseptically dispensed into sterile Erlenmeyer flasks. Finally, a soil infusion broth (SB) was created from a dried mixture of 80% soil (from INRAE Avignon, France) and 20% organic compost (wt/wt); 400 g of this mixture were autoclaved in 1 L of demineralized water at 121 °C for 60 minutes. After decanting overnight, the liquid fraction was clarified by paper filtration and subsequent centrifugation at 7,000 x g for 10 minutes, and the final supernatant was dispensed into flasks and sterilized by autoclaving at 121°C for 20 minutes.

An overnight preculture was prepared in 4 mL Luria-Bertani (LB medium (10g/L tryptone, 5g/L yeast extract, 10 g/L NaCl) at 37 °C. The preculture was centrifuged (7,000 x g, 6 min 10 °C), and the cell pellet was washed with 5 mL of one of our target media before being before being resuspended in 4 mL of the same fresh medium. A volume of these suspensions (1 to 3 mL) was used to inoculate 500 mL Erlenmeyer flasks containing 100mL volume the test broths to reach an initial optical density at 600 nm of 0.05. Flasks were kept under rotative agitation at 200 rpm at 37 °C until the optical density reached 0.5 (mid-exponential phase, typically within 2.5 to 3 h) for all media except the soil media cultures that were stop at the optical density of 0.25, which correspond for this media to the mid-exponential growth phase (typically around 5 h of culture for this broth).

Upon harvesting, cultures were centrifuged in 50 mL conical tubes (7,000 x g, 15 min, 4 °C), the cell pellets were then washed twice in 5mL of Cold PBS (Phosphate-buffered Saline: 0.137 M NaCl, 2.7mM KCl, 8 mM $Na_2HPO4$ and 2 mM $KH_2PO4$ pH 7.4). After the second wash, the pellet was resuspended in 1 mL of PBS to allow its transfer to a 1.5 mL microcentrifuge tube and was centrifuged again to remove the supernatant. The final pellet was stored at -80 °C, until analysis.

**Proteomics analysis of *B. cereus* pellets.**

Cell lysis was achieved through mechanical disruption. Pellets were resuspended in 200 µL of 100 mM $NH_4HCO_3$ and subjected to bead-beating in a Bullet Blender (Next Advance) with approximately 100 µL of 0.1-mm zirconia-silica beads for 4 minutes at 4 °C (speed 8). To maximize recovery, the beads were washed with an additional 200 µL of 100 mM $NH_4HCO_3$, and this supernatant was pooled with the initial cell lysate. The total protein concentration was then quantified using a bicinchoninic acid (BCA) assay (Thermo Fisher Scientific). For digestion, aliquots containing 300 µg of protein were denatured and reduced in 8 M urea with 5 mM dithiothreitol (DTT) by incubating at 60 °C for 30 minutes with agitation (850 rpm). After reduction, samples were diluted 10-fold with 100 mM $NH_4HCO_3$, supplemented with $CaCl_2$ to a final concentration of 1 mM, and digested with trypsin for 3 hours at 37 °C using a 1:50 enzyme-to-protein ratio. The resulting peptides were desalted using 50-mg C18 cartridges (Strata; Phenomenex) following supplier's instructions. Finally, peptide concentration was quantified again by BCA assay before the samples were arranged into blocks, randomized, and submitted for analysis by liquid chromatography-tandem mass spectrometry (LC-MS/MS). LC-MS/MS as follow, peptides were resuspended in water, and a total of 500 ng was loaded into a trap column (5 cm by 360-µm-outer-diameter by 150-µm-inner-diameter fused silica capillary tubing, Polymicro, Phoenix, AZ) packed with 3.6 µm Aeries C18 particles (Phenomenex, Torrance, CA). Separation was performed in a capillary column (70 cm by 360-µm OD by 75-µm ID) packed with 3-µm Jupiter C18 stationary phase (Phenomenex) with the following gradient: 1%–8% B solvent in 2 min, 8%–12% B in 18 min, 12%–30% B in 55 min, 30%–45% B in 22 min, 45%–95% B in 3 min, hold for 5 min in 95% B and 99%–1% B in 10 min. Solvent A was composed of 0.1% formic acid in water and solvent B, 0.1% formic acid in acetonitrile. Eluted peptides were analyzed online in a quadrupole-Orbitrap mass spectrometer (Q-Exactive Plus; Thermo Fisher Scientific, San Jose, CA). Tandem mass spectra were collected for the top 12 most intense ions. The Raw LC-MS/MS data is publicly available, on MassIVE under the identifier MSV000085696 (FTP at the following address: ftp://massive.ucsd.edu/MSV000085696/).

**Data analysis of *B. cereus*.**

Raw data was preprocessed with FragPipe v23.1 (with MSFragger V4.3 and IonQuant V1.11.11) searched against the UniProt proteome of *B. cereus* stain ATCC14579 (proteome ID: UP000001417 downloaded 10/10/2025, 5,240 sequence). The search fasta file included reverse sequences for decoy generation and FragPipe[21] generated list of common contaminants. The search was configured as semi-tryptic, allowing for up to two misscleavages. Precursor and fragment mass tolerances were set at 20 ppm.

Carbamidomethylation of cysteines (+57.02146 Da) were set as fixed modifications, while oxidation of methionine (+15.9949 Da) and protein N-terminal acetylation (+42.0106 Da) were set as variable modifications. The quantification was performed using the IonQuant MaxLFQ algorithm, and the match-between runs option was enabled.

Following the import of the dataset into a Romics_object, named "romics_proteins". The object was immediately duplicated into an object named "romics_unprocessed". Proteomics data processing was executed on "romics_proteins" as follows. Initially, zero values were converted to missing values. A filtering step was then applied to retain the proteins that were detected in at least 70% of the samples within any single experimental group. The remaining protein abundance values were log2-transformed followed by a sample-wise median centering. For data exploration, Principal Component Analysis (PCA was performed. To identify the differentially abundant proteins, different statistical tests were applied. A one-way ANOVA was used for global comparison across all groups. Pairwise comparisons were conducted using a two-sided T-test (assuming unequal variance), restricted to proteins with at least 70% completeness in one of the compared groups. In parallel, a GLM binomial test was used to identify proteins with significant presence/absence patterns across paired conditions. The steps of "romics_proteins" were extracted with the *romics_steps()* function. The object "romics_proteins" was then reinitialized under the name "romics_restored_original" using the function *resetRomicsObject()*. A pipeline was created from "romics_proteins" using the line of code

> *romics_pipeline<-createRomicsPipeline(romics_proteins)*

Then this pipeline was applied to the two objects "romics_restored_original" and "romics_unprocessed" and we verified that the data layer and statistics layers were identical bit-for-bit in the three objects.

The R markdown generated is provided on Zenodo (https://doi.org/10.5281/zenodo.21266809).

Below is the log of the "romics_protein$steps" layer at the end of the analysis

```
"date|Jun_04_2026_10:39:59|createRomicsObject"
"fun|createRomicsObject(data=data,metadata=metadata,IDs=IDs,main_factor='g
rowth_media',omics_type='proteomics',omics_information=omics_information)
"date|Jun_04_2026_10:39:59|romicsZeroToMissing"
"fun|romicsZeroToMissing(romics_object=romics_proteins)"
"date|Jun_04_2026_10:39:59|romicsFilterMissing"
"fun|romicsFilterMissing(romics_object=romics_proteins,percentage_complete
ness=70)"
"date|Jun_04_2026_10:39:59|log2transform"
"fun|log2transform(romics_object=romics_proteins)"
"date|Jun_04_2026_10:40:01|medianCenterSample"
"fun|medianCenterSample(romics_object=romics_proteins)"
"date|Jun_04_2026_10:40:03|romicsPCA"
"fun|romicsPCA(romics_object=romics_proteins)"
"date|Jun_04_2026_10:40:04|romicsMean"
"fun|romicsMean(romics_object=romics_proteins)"
"date|Jun_04_2026_10:40:04|romicsSd"
"fun|romicsSd(romics_object=romics_proteins)"
"date|Jun_04_2026_10:40:04|romicsZscores"
"fun|romicsZscores(romics_object=romics_proteins)"
"date|Jun_04_2026_10:40:06|romicsANOVA"
"fun|romicsANOVA(romics_object=romics_proteins)"
"date|Jun_04_2026_10:40:08|romicsTtest"
"fun|romicsTtest(romics_object=romics_proteins,alternative='two.sided',var
.equal=FALSE,percentage_completeness=70)"
"date|Jun_04_2026_10:40:28|romicsGlmBinomial"
"fun|romicsGlmBinomial(romics_object=romics_proteins)"
"date|Jun_04_2026_10:40:36|romicsChangeIDs"
"fun|romicsChangeIDs(romics_object=romics_proteins,newIDs='Gene')"
"date|Jun_04_2026_10:56:13|romicsChangeIDs"
"fun|romicsChangeIDs(romics_object=romics_proteins,newIDs='original')"
```

#### Human Tissue Microarray (TMA) generation

All human lung samples were obtained from the Biorepository for Investigation of Diseases of the Lung (BRINDL; https://brindl.urmc.rochester.edu) supported by the NHLBI LungMAP program and under protocol (dx.doi.org/10.17504/protocols.io.bjuxknxn). The use of the

BRINDL tissues as non-human subjects research is approved by the University of Rochester IRB (#RSRB00047606). Tissue microarrays (TMA) were constructed as previously described from formalin-fixed paraffin embedded donor lung tissue blocks (dx.doi.org/10.17504/protocols.io.kxygxejwdv8j/v3; dx.doi.org/10.17504/protocols.io.yxmvmmd46v3p/v3). Regions of interest were chosen in duplicate by a pathologist on hematoxylin and eosin (H&E) stained slides, with the corresponding area of the paraffin blocks retrieved by 3.0 mm biopsy punch and vertically hand-placed and paraffin-embedded in a metal-based mold according to a TMA construction map. Serial sections were generated from the constructed TMA including an H&E section to validate accurate representation and placement of all the cores according to the TMA slide map (mirror image of TMA construction map). Donor demographics are reported in Supplemental Table 1.

**Multiplexed Immunohistochemistry.** TMA sections were assessed by multiplexed immunohistochemistry using the Phenocycler-Fusion CODEX system (Akoya Biosciences, Marlborough, MA) and our published protocol (dx.doi.org/10.17504/protocols.io.6qpvr38dpvmk/v3). Candidate protein targets were selected based on relative abundance, and anticipated cell type specificity based on our prior transcriptomic and proteomics studies. Selected antibodies were either purchased with oligonucleotide tags (Akoya Biosciences) or conjugated to oligonucleotide barcodes on site as described (dx.doi.org/10.17504/protocols.io.3fugjnw). The antibodies were then developed into a fifty-antibody panel with concentrations, channels and run combinations optimized. Validation of Ab staining was performed as previously described for lung OMAP panels **dx.doi.org/10.17504/protocols.io.rm7vzkm14vx1/v1**. The sources, clones and concentrations of antibodies used in the Romics test case are reported in Supplemental Table 2.

**CODEX pre-processing** CODEX data were preprocessed using Mesmer DeepCell (0.12.10) in python (3.8.10). Masks were generated using Mesmer Deep Cell with DAPI as the nuclear marker and ATP1A1 as the membrane marker. To account for limitations in RAM, the original image was divided into tiles. Each tile and mask were run through region_props_table() in scikit-image (0.21.0) to generate data tables to return the average intensity value within each cell mask for each marker. The results were stacked and collated into a single AnnData object using anndata (0.9.2) and scanpy (1.9.8). Signal intensity and obs metadata tables from this object were converted to csv files and used in RomicsProcessor.

**MALDI-MSI Lipidomics.** Kidney biopsies were procured as part of the kidney precision medicine project (KPMP) from 4 subjects, control tissue (n = 3) and acute kidney injury (AKI = 3) (Supplementary Table 3). The fresh-frozen renal tissues were sectioned at 7 µm using a cryostat set at -18 °C and thaw-mounted on indium tin oxide-coated glass microscopy slides (Delta Technologies). The 2,5-dihydroxybenzoic acid matrix was applied to the slides (40 mg/ml in 70% MeOH:$H_2O$) using an automated HTX M5 sprayer with nozzle temperature 75 °C, a flow rate of 0.05 ml/min, 10 passes, nitrogen pressure of 10 psi, 40 mm nozzle height, and track spacing of 3 mm. MALDI-MSI was performed using a Bruker 12-Tesla (12T) Fourier-transform ion cyclotron resonance (FTICR) mass spectrometer equipped with a SmartBeam II laser source (355 nm, 2 kHz) in positive ion mode using a 25 µm step-size. FTICR-MS was operated to collect 240 – 1100 m/z with a mass resolving power of ~160,000 at 400 m/z. Ion transfer parameters were optimized for the detection of 600-900 m/z ions.

**Histology Staining.** After MALDI-MSI, the matrix is removed using 70 % methanol for 1 min, and periodic acid shift staining is performed. The slides are rinsed four times in water and placed in periodic acid solution for 5 min. Slides were rinsed with water four times and placed in Schiff's reagent for 5 min, followed by an additional rinse in water for 5 min. Then they were counterstained in Hematoxylin for 3 min, dipped into TRIS buffer saline (TBS), washed with water four times, dehydrated for 2 min in 100% ethanol twice, and xylene twice. Each tissue section was coverslipped and imaged using an Aperio ScanScope XT at 20x magnification. The tubular interstitium and glomeruli of each sample were manually annotated in QuPath (v-0.3.2) before importation into Bruker's SCiLS software using the QuPath to SCiLS extension.

**MALDI Data Processing.** Bruker's SCiLS software was used to convert each dataset into the vendor-neutral IMZML format, followed by uploading to the open-source online platform, METASPACE, for annotation against the SwissLipids and an internal LC-MS/MS database created using kidney tissues. The molecular annotations from SwissLipids FDR < 20% were concatenated with the molecular annotations from LC-MS/MS database and imported into SCiLS for downstream analysis. SCiLS data files containing the molecular annotations, ion abundances, and histology segmentations were exported to R using the SCiLS API, and data processing was performed in *ROmicsProcessor*.

| HuBMAP ID | Donor ID | Age (yrs) | Sex | Race/ Ethnicity | Cause of Death | ARDS Status | Histopathological Findings | Past Medical History | Wt (kg) | BMI | WIT (hrs) | CIT (hrs) |
|---|---|---|---|---|---|---|---|---|---|---|---|---|
| HBM443. VFRD.453 | D231 | 33 | F | Black/AA | Anoxic Brain Injury | neg | Normal lung structure; Patchy mild periairway fibrosis, increased smooth muscle; Focal prominence bronchial goblet cells; Increased alveolar macrophages; Patchy mild mixed inflammation; Mild large pulmonary artery branch intimal hyperplasia, Scattered thrombi (microvascular) | Stable, chronic ventilation due to brain injury; Renal calculi | 104 | 36 | 0 | 29 |
| HBM943. SCQQ.877 | D239 | 37 | M | Black/AA | Stroke | mild | Largely normal alveolar structure; Mild bronchiectasis and goblet cell hyperplasia; Foci of bronchiolar metaplasia; Marked occlusive intimal hyperplasia of bronchial vessels; Medial hypertrophy and intimal hyperplasia of large and medium-sized veins; Scattered hemosiderin laden macrophages | Hypertension, obesity; Tox screen + cannabinoids; | 151 | 45 | 0 | 19 |
| HBM643. KCCK.866 | D341 | 56.8 | F | Hispanic | Stroke | neg | Normal alveolar structure; Medial hypertrophy and focal intimal fibrosis of large pulmonary arteries; Acute and calcified arterial thrombi as well as rare microvascular thrombi; Intimal pulmonary vein fibrosis; Bronchiolar dilation with peri airway anthracitic pigment and focal mild mucostasis | Hypertension, polycystic kidney disease, breast cancer > 5 yrs prior | 48 | 20 | 0 | 34 |

| | | | | | | | | | | | | |
|---|---|---|---|---|---|---|---|---|---|---|---|---|
| HBM932. JNVS.672 | D346 | 59 | M | White | Stroke | mild | Normal alveolar and airway structure; Hypertensive vascular changes with medial hypertrophy, intimal fibrosis | Hypertension | 70 | 23 | 0 | 26 |
| HBM565.TRST.423 | D438 | 24 | F | Black/AA | Brain Injury | neg | Normal lung structure; Minimal patchy acute airway inflammation, some hemorrhage | Seasonal Allergies; Every other day marijuana | 106 | 40 | 0 | 39 |
| HBM566.WFPK.939 | D460 | 20 | M | Black/AA | Brain Injury | neg | Normal lung structure; Patchy acute inflammation; Extensive acute alveolar hemorrhage, scattered hemosiderin-laden macrophages | None | 91 | 27 | 0 | 27 |

**Supplementary Table 1.** Lung TMA Donor Demographics and Histological Assessment

| UniProt # | HGNC_ID | Protein | Target (Level if applicable) | Vendor | Host (clonality) | Isotype | Clone Id | RRID | Dilution Factor |
|---|---|---|---|---|---|---|---|---|---|
| | | DAPI Stain | DNA | | | | | | |
| P62736 | HGNC:130 | ACTA2 | CT.Muscle (L1) | Akoya | Mouse (m) | IgG2a | 1A4 | AB_2936084 | 200 |
| Q15109 | HGNC:320 | AGER | AT1 (L4) | Abcam | Rabbit (m) | IgG | EPR21171 | AB_2884897 | 100 |
| P54709 | HGNC:806 | ATP1A1 | Cell segmentation | Abcam | Rabbit (m) | IgG | EP1845Y | AB_2890241 | 100 |
| Q03135 | HGNC:1527 | CAV1 | Endothelial (Superclass), AT1 (L4) | Akoya | Rabbit (m) | IgG | D46G3 | AB_3508120 | 200 |
| P07766 | HGNC:1674 | CD3E | T (L3) | Akoya | Rabbit (m) | IgG | EP449E | AB_2936080 | 200 |
| P01730 | HGNC:1678 | CD4 | CD4.T (L5) | Akoya | Rabbit (m) | IgG | EPR6855 | AB_2915936 | 200 |
| P34810 | HGNC:1693 | CD68 | Imm.Myeloid (L1), Macrophage (L2) | Akoya | Mouse (m) | IgG1 | KP1 | AB_2935894 | 200 |
| P01732 | HGNC:1706 | CD8A | CD8.T cell (L5) | Akoya | Mouse (m) | IgG1 | C8/144B | AB_2915960 | 200 |
| P12830 | HGNC:1748 | CDH1 | Epithelial (Superclass) | Akoya | Mouse (m) | IgG1 | 4A2C7 | AB_2895057 | 200 |
| Q05707 | HGNC:2202 | COL14A1 | Extracellular matrix | Cell Signaling | Rabbit (m) | IgG | E5W8S | AB_3696891 | 200 |
| P02452 | HGNC:2197 | COL1A1 | Extracellular matrix | Abcam | Mouse (m) | IgG3 | 3G3 | AB_2081873 | 100 |
| P02462 | HGNC:2202 | COL4A1 | Extracellular matrix | Akoya | Rabbit (m) | IgG | EPR20966 | AB_2927676 | 200 |
| P12109 | HGNC:2211 | COL6A1 | Extracellular matrix | Abcam | Rabbit (m) | IgG | EPR17072 | AB_2847919 | 200 |

| | | | | | | | | | |
|---|---|---|---|---|---|---|---|---|---|
| Q9Y6C2 | HGNC:19880 | EMILIN1 | AlvFB-2 (L5), Extracellular matrix | Abcam | Rabbit (m) | IgG | EPR14678 | AB_3696069 | 500 |
| P02751 | HGNC:3778 | FN1 | AlvFB (L1), Extracellular matrix | Abcam | Rabbit (m) | IgG | EPR23110-46 | AB_2927764 | 1000 |
| Q12951 | HGNC:3815 | FOXI1 | Ionocytes (L5) (nuclear) | LSBio | Mouse (m) | IgG1 | OTI1D4 | AB_3696022 | 100 |
| Q92949 | HGNC:3816 | FOXJ1 | Multiciliated (L4) (nuclear) | ThermoFischer | Mouse (m) | IgG1 | 2A5 | AB_1548835 | 500 |
| Q9BZS1 | HGNC:6106 | FOXP3 | CD4.T (L5) | Akoya | Mouse (m) | IgG1 | 236A/E7 | AB_2927679 | 200 |
| P42261 | HGNC:4571 | GRIA1 | AlvFB (L4) (suboptimal stain) | Abcam | Rabbit (m) | IgG | EPR19522 | AB_3696021 | 400 |
| P07492 | HGNC:4605 | GRP | PNEC (L5) | LSBio | Rabbit (p) | IgG | (polyclonal) | AB_3096183 | 500 |
| P15428 | HGNC:5154 | HPGD | CAP2 (L5) | Abcam | Rabbit (m) | IgG | EPR14332 | AB_2861359 | 500 |
| P53708 | HGNC:6144 | ITGA8 | AlvFB (L4) (suboptimal stain) | ThermoFischer | Mouse (m) | IgG1 | CL7304 | AB_2787085 | 400 |
| P02533 | HGNC:6416 | KRT14 | SMG Duct Epith, MEC (L5) | Akoya | Rabbit (p) | IgG | Poly19053 | AB_3095339 | 200 |
| Q04695 | HGNC:6427 | KRT17 | Basal (L5), MEC (L5) | Abcam | Rabbit (m) | IgG | EP1623 | AB_3696892 | 400 |
| P13647 | HGNC:6442 | KRT5 | Basal (L5) | Akoya | Rabbit (m) | IgG | EP1601Y | AB_3083458 | 200 |
| P05787 | HGNC:6446 | KRT8 | Epithelial (Superclass), AT2 (L4) | BioLegend | Mouse (m) | IgG2a | 1E8 | AB_2616821 | 400 |
| P02788 | HGNC:6720 | LTF | Serous (L5) | Proteintech | Rabbit (p) | IgG | (polyclonal) | AB_2139227 | 500 |
| Q9Y5Y7 | HGNC:14687 | LYVE1 | Endo.Lymph (L1), LEC (L4) | R&D Systems | Goat (p) | IgG | (polyclonal) | AB_355144 | 100 |
| P50221 | HGNC:7013 | MEOX1 | Endothelial subset (nuclear) | LSBio | Mouse (m) | IgG1 | OTI5B11 | AB_3696022 | 400 |
| Q13361 | HGNC:29673 | MFAP5 | AdvFB (L2), NeurFB (L2) | Abcam | Rabbit (m) | IgG | EPR17581 | AB_3696411 | 500 |
| P46013 | HGNC:7107 | MKI67 | Proliferation | Akoya | Mouse (m) | IgG1 | B56 | AB_2895046 | 200 |
| P05164 | HGNC:7218 | MPO | Neutrophil (L4), Macrophage (L2) | Akoya | Rabbit (m) | IgG | E1E7I | AB_2927678 | 200 |
| P11836 | HGNC:7315 | MS4A1 | B (L5) | Akoya | Mouse (m) | IgG2a | L26 | AB_2915939 | 200 |
| P98088 | HGNC:7515 | MUC5AC | Goblet (L5) | Abcam | Mouse (m) | IgG1 | 45M1 | AB_3083499 | 100 |
| Q9HC84 | HGNC:7516 | MUC5B | Goblet (L5) | Abcam | Rabbit (p) | IgG | (polyclonal) | AB_10712492 | 75 |

| | | | | | | | | | |
|---|---|---|---|---|---|---|---|---|---|
| P04264, P35908, P12035, P19013, P13647, P02538, P04259, P48668, P08729, P05787, P13645, P02533, P19012, P08779, P08727 | HGNC:6412, HGNC:6439, HGNC:6440, HGNC:6441, HGNC:6442, HGNC:6443, HGNC:6444, HGNC:20406, HGNC:6445, HGNC:6446, HGNC:6413, HGNC:6416, HGNC:6421, HGNC:6423, HGNC:6436 | (PANCK) KRT1, KRT2, KRT3, KRT4, KRT5, KRT6A, KRT6B, KRT7, KRT8, KRT10, KRT14, KRT15, KRT16, KRT19 | Epithelial (Superclass) | Akoya | Mouse (m) | IgG1 | AE-1/3 | AB_3083456 | 200 |
| Q86YL7 | HGNC:29602 | PDPN | AT1 (L4), Endo.Lymph (L1), LEC (L4) | Akoya | Mouse (m) | IgG2a | NC-08 | AB_3082979 | 200 |
| P16284 | HGNC:8823 | PECAM1 | Endothelial (Superclass) | Akoya | Rabbit (m) | IgG | EP3095 | AB_2915935 | 200 |
| P02776 | HGNC:8861 | PF4 | Platelet | PeproTech | Rabbit (p) | IgG | (polyclonal) | AB_147868 | 200 |
| Q9H5I5 | HGNC:26270 | PIEZO2 | AlvFB-2 (L5) | Novus | Rabbit (p) | IgG | (polyclonal) | AB_11005294 | 100 |
| Q92786 | HGNC:9459 | PROX1 | Endo.Lymph (L1) | Abcam | Rabbit (m) | IgG | EPR19273 | AB_2894898 | 200 |
| P08575 | HGNC:9666 | PTPRC | Immune (Superclass) | Akoya | Rabbit (m) | IgG | D9M8I | AB_2915946 | 200 |
| O95171 | HGNC:10573 | SCEL | AT1 (L4) | Abcepta | Rabbit (p) | IgG | (polyclonal) | AB_10818433 | 100 |
| P11684 | HGNC:12523 | SCGB1A1 | Aw.Secretory (L2) | R&D | Rat (m) | IgG1 | 394324 | AB_2183286 | 400 |
| Q96PL1 | HGNC:18391 | SCGB3A2 | BrAS (L2) | Abcam | Rabbit (m) | IgG | EPR11463 | AB_3083498 | 400 |
| P07988 | HGNC:10801 | SFTPB | Basal.SFTPB (L4), Club.SFTPB (L4), BrAS (L2), AT2 (L4) | Invitrogen | Rabbit (p) | IgG | (polyclonal) | AB_2609628 | 50 |
| P11686 | HGNC:10802 | SFTPC | AT2 (L4) | Invitrogen | Rabbit (p) | IgG | (polyclonal) | AB_2717696 | 500 |
| Q9H3D4 | HGNC:15979 | TP63 | Basal (L5), MEC (L5) | Abcam | Rabbit (m) | IgG | EPR5701 | AB_3083495 | 100 |

| | | | | | | | | | |
|---|---|---|---|---|---|---|---|---|---|
| Q15661 | HGNC:12019 | TPSAB1 | Mast (L2) | Abcam | Mouse (m) | IgG1 | AA1 | AB_303023 | 1000 |
| Q13509 | HGNC:20772 | TUBB3 | Neural (L2) | R&D | Mouse (m) | IgG2a | TUJ-1 | AB_357520 | 400 |
| P19544 | HGNC:12796 | WT1 | Mesothelial (L5), SubpleurFB (L5) | Novus | Mouse (m) | IgG1 | 6F-H2 | AB_905863 | 200 |

**Supplementary Table 2. Antibodies Used in Multiplexed Immunofluorescence (MxIF)** Target proteins and antibody clone IDs, commercial source and dilution factors. (RRID at www.antibodyregistry.org; Clonality: (m) monoclonal, (p) polyclonal)

| Participant ID | Disease Category | Age | Gender | Race |
|---|---|---|---|---|
| 32-10346 | AKI | 44 | Male | Black |
| 32-10456 | AKI | 40 | Male | Black |
| 30-11081 | AKI | 73 | Female | White |
| 164-18 | HRT | 46 | Male | White |
| 077 | HRT | 52 | Male | White |

**Supplementary Table 3.** Kidney biopsy sample metadata
